# Improved crystallization and diffraction quality of *Mycobacterium tuberculosis* OmamC/Rv1363c upon heat treatment

**DOI:** 10.64898/2026.04.30.722021

**Authors:** Mikko J Hynönen, Rajaram Venkatesan

## Abstract

*Mycobacterium tuberculosis* (Mtb), the causative agent of tuberculosis, can use host derived lipids as carbon and energy source for survival. Mammalian cell entry (Mce) associated membrane (Mam) proteins are important for the stability of lipid importing Mce complexes. Mtb has five homologs of Mam proteins referred as orphaned Mam (OmamA-E) proteins. A recent study suggested that OmamC (Rv1363c) is essential for the storage and utilization of lipids under starvation in Mtb. To understand the structure and interactions of OmamC, we generated a truncated soluble variant of OmamC (OmamC_129-261_). Here, we report on the challenges encountered during the crystallization and structure determination of OmamC_129-261_ and the strategies applied to overcome them. Despite the AlphaFold2 predicted model proving an initial molecular replacement solution, experimental phasing was necessary to determine the structure of OmamC_129-261_. Heat treatment of protein prior to crystallization setup removed partially unfolded protein present and played a critical role in enhancing the reproducibility and diffraction quality of OmamC_129-261_ crystals. Although reported earlier, it is not a widely used method. It is worth to try this method, especially, when faced with poor reproducibility and diffraction of crystals.

## Introduction

*Mycobacterium tuberculosis* (*Mtb*), the causative agent of tuberculosis (TB) is one of the cleverest pathogens that has co-evolved with human for centuries and leads to more than a million deaths per year^1^. A key mechanism that *Mtb* uses to survive in the host cell is by using the host-derived lipids as carbon sources^2^. Mammalian cell entry (Mce) complexes Mce1-4 in *Mtb* have shown to be importers of such lipids^3–5^. The operons encoding Mce complexes also encodes Mce-associated membrane (Mam) proteins which are suggested to be stabilizing proteins^5,6^. Homologs of Mam proteins have been identified in *Mtb* and related organisms which are encoded elsewhere in the genome and therefore named as orphan Mam (Omam) proteins^7^. *Mtb* has five Omam proteins, OmamA (Rv0199), OmamB (Rv0200), OmamC (Rv1363c), OmamD (Rv1362c), and OmamE (Rv2390c). Of this, OmamA has shown to be associated with Mce1 and Mce4 complexes^6,8^. However, the role of other Omam proteins have not been understood. In a recent study, OmamC has been shown to be important for the *Mtb* to stay in hibernation through the utilization of stored fatty acids. In an effort to understand the role of OmamC, structural studies were initiated on the soluble variant of OmamC, OmamC_129-261_ with the variable N-terminal region and membrane spanning helix removed. During the crystallization and structure determination process of OmamC_129-261_, several interesting observations were made which are reported here. Especially, the diffraction quality of the crystals improved by pre-heating the sample before setting up crystallization. The phases had to be determined experimentally using SeMet SAD phasing method, despite the AlphaFold2^9^ model providing an initial solution in molecular replacement method.

## Results and discussion

### Deletion of 128 amino acids from the N-terminus of OmamC results in a soluble OmamC_129-261_ variant

OmamC, like other Mam/Omam proteins have four conserved regions^10^ i) a variable N-terminal region, ii) a long alpha helical region including transmembrane (TM) region iii) two helices linked by short loops and iv) four beta-strands. In OmamC, the N-terminal variable region has 106 residues. Despite optimization and use of different detergents for purification, a monodisperse population of OmamC could not be obtained. To improve solubility and stability, the variable N-terminal region and part of the helix predicted to be spanning the membrane was removed, creating OmamC_129-261_ variant. This variant could be purified without the use of any detergent.

OmamC_129-261_ was purified using immobilised metal affinity chromatography (IMAC) followed by size exclusion chromatography (SEC). The SEC profile in a HiLoad 16/60 Superdex 200 prep grade 120 ml column showed two peaks, one at 56 ml and another at 92 ml (**Fig. 1A**). Analysis of the peak fractions showed that both the peaks contained OmamC_129-261_. However, peak1 had more impurities. Therefore, only fractions from peak2 were pooled, concentrated and used for further characterization. Circular dichroism (CD) spectroscopy measurements showed that OmamC_129-261_ is well folded (**Fig. 1B**), and a thermal stability analysis using nano differential scale fluorimetry (nanoDSF) also suggested OmamC_129-261_ to be a well-folded protein with a *T*_m_ of 45 °C, Δ*G*_u_ of 11.7 8 kcal mol^-1^, and a Δ*H*_m_ of 186 kcal °C^-1^ (**Fig. 1C**). Glycerol is commonly used to improve solubility of membrane related proteins, and so the purification was repeated in the buffer containing 5% glycerol. Although the SEC profile was similar to that purified in the absence of glycerol, CD analysis of peak2 fractions showed that the protein is not well-folded (**Fig. 1B**). Furthermore, T_m_ calculated from nanoDSF was unusually high with 72 °C but with very low Δ*G*_u_ (3.8 kcal mol^-1^) and ΔH_m_ (27.8 kcal °C^-1^) values suggesting that the protein is unfolded, or at best poorly folded (**Fig. 1C and Table 1**). This is also evident from the lack of appropriately shaped curve in the nanoDSF measurement. Therefore, further characterization was carried out with the protein purified with the buffers not containing glycerol.

**Table 1.** Thermal stability calculated from nanoDSF measurements using Moltenprot webservice.

| Variant | $T_m$, °C | $\Delta G_u$, kcal mol<sup>-1</sup> | $\Delta H_m$, kcal °C<sup>-1</sup> |
| --- | --- | --- | --- |
| OmamC<sub>129-261</sub> Peak 1 | 62.3 | 1.75 | 15.8 |
| OmamC<sub>129-261</sub> Peak 2 | 45.0 | 11.7 | 186 |
| OmamC<sub>129-261</sub> heat treated | 46.3 | 10.0 | 150 |
| OmamC<sub>129-261</sub>SeMet peak 1.5 | 45.4 | 6.9 | 107 |
| OmamC<sub>129-261</sub>SeMet peak 2 | 44.8 | 7.6 | 122 |
| OmamC<sub>129-261</sub> with glycerol | 72 | 3.8 | 27.8 |

**Table 2.** Data collection and refinement statistics of the OmamC_129-261_ datasets.

|  | **OmamC<sub>129-261</sub>** | **OmamC<sub>129-126SeMet</sub>** |
| --- | --- | --- |
| **Data collection parameters** |  |  |
| **Beamline** | I04, Diamond Light Source, UK | I04, Diamond Light Source, UK |
| **Wavelength, Å (eV)** | 1.06471 (11,645) | 0.9795 (12,658) |
| **Exposure (seconds)** | 0.0087 | 0.0123 |
| **Dose (MGy)** | 3 | 3 |
| **Total collected images** | 1200 | 1200 |
| **Oscillation (°)** | 0.15 | 0.15 |
| **Data collection statistics (\* Values in parenthesis correspond to highest resolution shell)** |  |  |
| **Space group** | C121 | P61 2 2 |
| **Cell dimension**<br>**a, b, c (Å)** | 71.12, 111.89, 75.24 | 65.85, 65.85, 434.69 |
| **Cell dimensions**<br>**$\alpha$, $\beta$, $\gamma$ (°)** | 90, 117.83, 90 | 90, 90, 120 |
| **Resolution, (Å)** | 66.54-2.3 (2.38-2.3)* | 72.45-3.14 (3.36-3.14) |
| **# of unique reflections,**<br>**Overall (outer shell)** | 22461 (1706) | 10867 (1871) |
| **Mean $\langle I/\sigma(I) \rangle$** | 6.4 (1.0) | 21.1 (5.7) |
| **Redundancy** | 3.4 (2.7) | 19 (20) |
| **Wilson B-factor, (Å<sup>2</sup>)** | 50.2 | 69.9 |
| **CC<sub>1/2</sub> (%)** | 99.5 (54.8) | 99.8 (98.1) |
| **CC<sub>1/2</sub> Anomalous (%)** | - | 100 (100) |
| **Redundancy**<sub>Anomalous</sub> | - | 10.5 (10.5) |
| **Refinement statistics** |  |  |
| **Refinement program** | phenix.refine, PHENIX<br>1.19.2_4158 |  |
| **R<sub>work</sub> (%)** | 22.81 |  |
| **R<sub>free</sub> (%)** | 25,27 |  |
| **# of atoms** | 3596 |  |
| **# of waters** | 78 |  |
| **Ramachandran plot (%)** |  |  |
| **Favoured/Allowed/Outliers** | 96.83/3.17/0.00 |  |
| **Rotamer outlier (%)** | 0.00 |  |
| **RMSD** |  |  |
| **bond lengths, (Å)** | 0.08 |  |
| **bond angles, (°)** | 0.21 |  |
| **Clash score** | 2 |  |
| **MolProbity score** | 1.2 |  |
| **Chain breaks, missing amino acid residues #** | B: 230 – 233<br>C: 228 – 235 |  |
| **Deposition** |  |  |
| **PDB id** | 9R33 |  |
| **Extended PDB id** | pdb_00009R33 |  |

**Fig 1.**
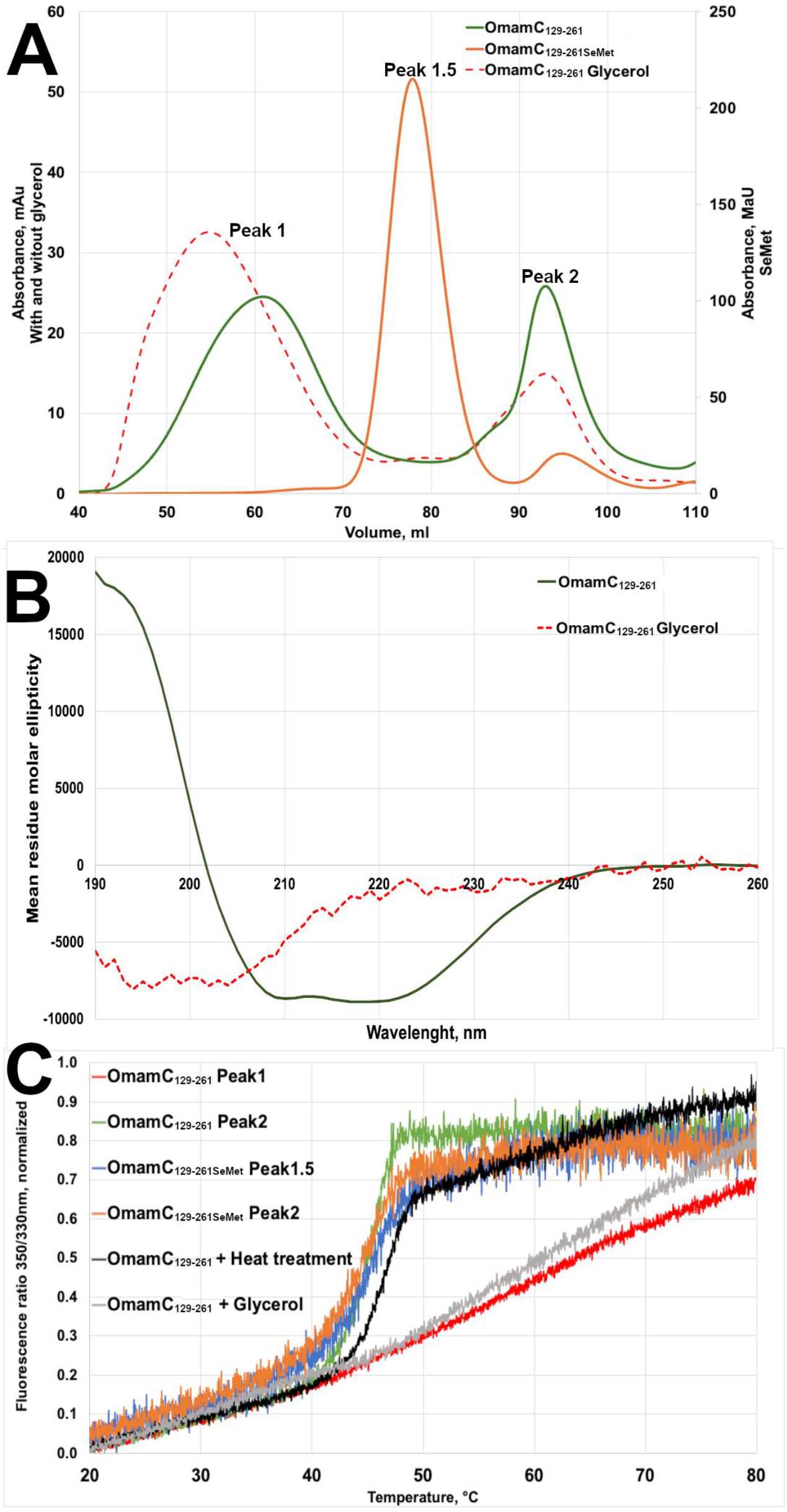
SEC and biophysical analysis of OmamC_129-261_. **A)** Comparison of SEC profile of OmamC_129-261_ purified under different conditions. OmamC_129-261_ (green line) and OmamC_129-261_ in the presence of glycerol (dotted red line) produced two peaks referred as peak 1 and peak 2. Seleno methionine labelled variant also produced two peaks (orange line), one referred as peak 1.5 as it lies between the peaks 1 and 2 of unlabelled OmamC_129-261_) and another as peak 2. **B)** CD spectrum of OmamC_129-261_ with (red dotted line) and without glycerol (green solid line) indicating significant loss of secondary structure elements in the presence of glycerol. **C)** nanoDSF fluorescence ratio curve of OmamC_129-261_ purified in different conditions. In the presence of glycerol OmamC_129-261_ is not well folded consistent with the CD profile. The parameters calculated form nanoDSF are summarized in **Table 1**.

### Initial crystallization of OmamC_129-261_ and structure determination trials by molecular replacement

The first crystals of OmamC_129-261_ appeared in the sub-well condition H03 of MIDASplus (MD1-106) screen which contained 30% glycerolethoxylate and 3% imine polymer in 100 mM Tris buffer at pH 8.5 (**Fig. 2A**). Also, the crystals were observed only in the plates incubated at 6 °C. From one of these crystals a dataset was collected to a resolution of 2.9 Å. The crystals belong to the space group C121. Matthew’s coefficient analysis indicated 3 molecules in the asymmetric unit. The closest available structural homolog is VirB8 (PDB id: 2CC3)^11^ with a sequence identity of about 14%. During this time, AlphaFold2 database was released which had a predicted structure for *Mtb* OmamC. This AlphaFold2 predicted model was used as the search model in molecular replacement trials in PHASER ^12^. This resulted in the placement of three molecules in the asymmetric unit with a translation function Z-score (TFZ) of 11.2 and log-likelihood gain (LLG) of 201 indicating that the model had been placed in the correct orientation and position in the new unit cell. However, further model building and refinement did not lead to an improved *R*_free_ or electron density. While the electron density corresponding to some parts of the structure, especially the N-terminal helix was good, the density connecting the different β-strands was very poor or not present at all. Various strategies used to improve the phases from this starting model did not reduce the *R*_free_ beyond 0.41.

**Fig 2.**
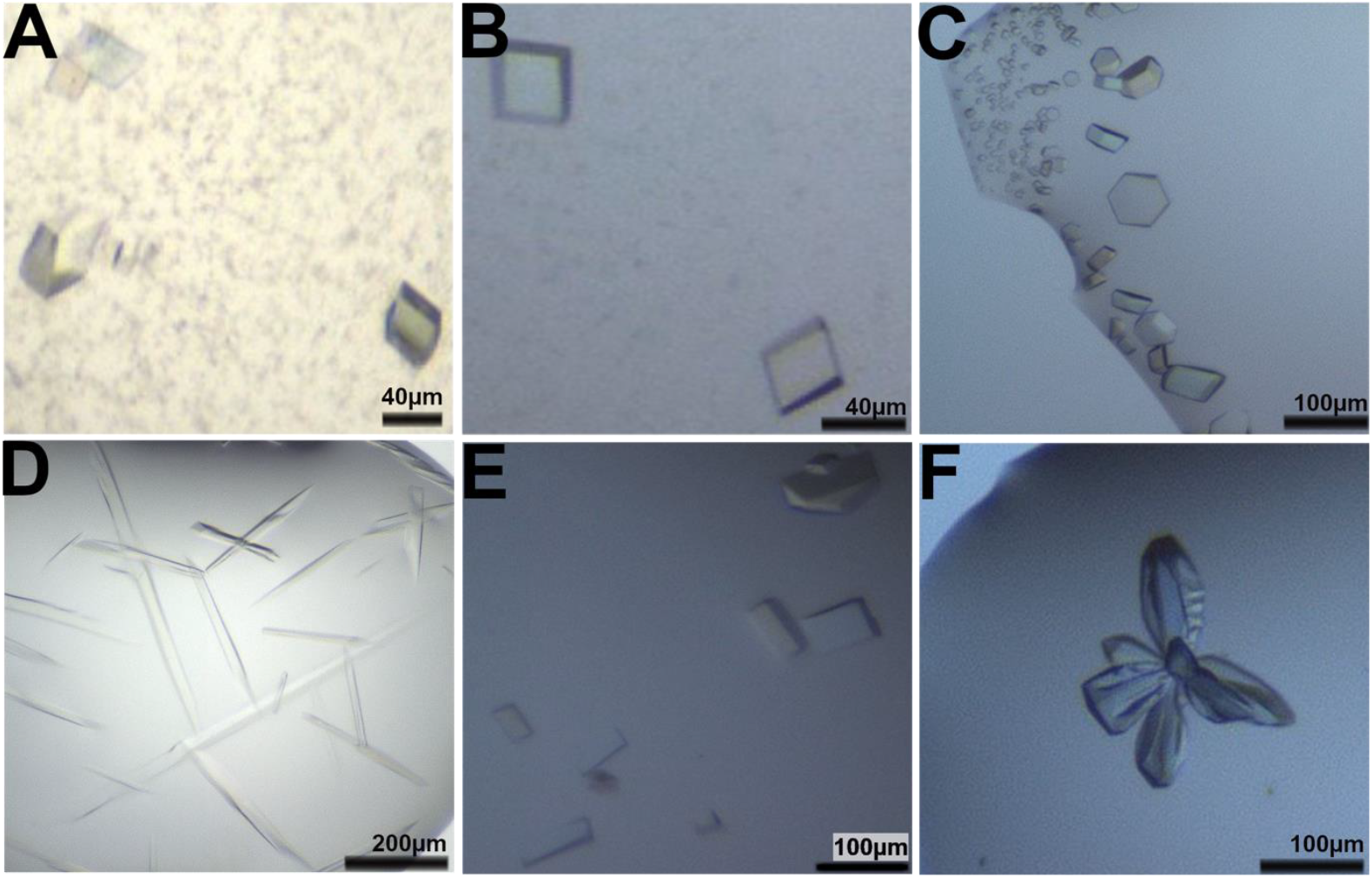
OmamC_129-261_ crystals produced from different crystallization trials. **A**) OmamC_129-261_ crystals obtained from MIDASplus sub-well H03 condition which diffracted to about 2.9 Å resolution **B-C**) Crystal of OmamC_129-261_ produced after heat treatment in different crystal morphologies. Crystals in Panel B diffracted to 2.3 Å. **D)** Crystals of OmamC_129-261SeMet_ from peak 1.5 with 1.5 M NaCl in the reservoir solution (alternate reservoir method). **E-F**) Crystals of OmamC_129-261SeMet_ after heat treatment improved diffraction to 2.58 Å resolution.

### Reproducibility of crystallization and improving the diffraction

As the phases could not be improved, trials to determine the structure by experimental phasing were initiated. However, reproducibility of the crystals was a challenge. Crystals appeared only if the same solution which gave crystals was used in the crystallization. Crystallization solutions prepared with the same formulation did not produce crystals. One key observation was that if a particular old stock of imine polymer borrowed from a neighbouring laboratory was used, the crystals were reproducible to some extent. The freshly bought imine polymer had a warning that it is light sensitive. Possibly, the old imine polymer had undergone some changes with light exposure over time which promoted the crystallization of OmamC_129-261_. The older imine polymer was also more viscous compared to the new imine polymer solution. Trials with increased concentration of new imine polymer still did not produce any crystals.

Importantly, the recreation of crystallization condition was observed to be extremely poor. Creating a copy of the initial crystallization condition using reagent from the lab, had low chance of creating nucleation. Therefore, once crystallization condition was observed, the mother liquid was stored to be use as the drop precipitant solution, while reservoir solution was prepared separately with the robot. Theoretically, both drop and reservoir solution should be of perfect match, but salvaging the mother liquid from previously observed crystallization plates immensely improved the frequency of observed crystals. However, the overall quality of the diffraction of these crystals did not improve.

A study by Pusey et al. (2005) suggested heat treatment of the purified protein sample as a way to improve crystal diffraction, by removing the partially unfolded proteins from solution.

Another case study also suggested the application of heat treatment to improve diffraction in protein crystals^13^. A similar methodology was attempted for OmamC_129-261_. nanoDSF analysis suggested that the thermal unfolding starts around 35 °C (**Fig. 1C**). Therefore, OmamC_129-261_ was incubated at various temperatures (30-45 °C) for 15 mins and the nanoDSF profile was tested after removing the precipitation. 35 °C was found to be optimal such that the resulting protein was still well-folded with a T_m_ similar to that observed for untreated OmamC_129-261_. The concentration of the clarified protein solution after heat treatment has reduced by about 30-50% from the original concentration. The nanoDSF profile indicated that the protein is intact with a similar *T*_m_ of about 46 °C, and further analysis calculated Δ*G*_u_ of 11.7 kcal mol^-1^, and a Δ*H*_m_ of 186 kcal °C ^-1^ (**Fig. 1C and Table 1**). Despite the lower starting concentration, the quality of the resulting crystals improved significantly in terms of size, frequency of crystallization as well as diffraction quality. However, the crystallization still required the use of original screen solution from MIDASplus H03 condition in the drop. This solution was salvaged from the crystallization wells which produced crystals earlier. Combining these two approaches, the diffraction of the crystals improved to 2.3 Å resolution (**Fig. 2B**). Despite these improvements in the crystallization, heavy atom soaking did not help to get a derivative suitable for phase determination. Therefore, the focus shifted towards producing seleno methionine (SeMet)-labelled protein and crystals.

### Purification and crystallization of OmamC_129-261SeMet_

SeMet-labelled OmamC_129-261_ (OmamC_129-261SeMet_) expressed in the B834 strain in a minimal media supplemented with SeMet was purified using a similar protocol as the native OmamC_129-261_. The incorporation of SeMet was confirmed by mass spectrometry. DTT was used in the purification buffer to minimize the oxidation of SeMet. In SEC, there was an additional peak between the peak 1 and peak 2 observed in OmamC_129-261_ (**Fig. 1A**), which was named as peak 1.5, which had a retention volume of about 78 ml. Peak 1.5 did not appear for native OmamC_129-261_ even when DTT was used in the buffer. The peak 1.5 behaved better with lower polydispersity and better unfolding curve in nanoDSF when compared to the peak 1 of native OmamC_129-261_ (**Fig. 1C and Table 1**).

The crystallization of OmamC_129-261SeMet_ was equally tricky. In the initial screening, no crystals appeared. However, an alternative reservoir method^14^ using 1.5 M NaCl in the reservoir produced crystals in multiple conditions, previously not observed in the classical screening.

Although further optimization with additive screening produced big (600 µm) crystals, they did not diffract beyond 5 Å. Therefore, the heat-treatment approach used for improving the quality of native OmamC_129-261_ crystals was applied also to OmamC_129-261SeMet_. Heat treated OmamC_129-261SeMet_ behaved in a similar way and produced crystals in the same condition as that of native OmamC_129-261_. These crystals diffracted to 2.58 Å, a significant improvement from the previously tested crystals. Anomalous signal was up to 3.14 Å.

### Structure of OmamC_126-261_

Despite being crystallized in the same condition as native OmamC_129-261_, the SeMet-labelled protein crystallized in a different space group (P6122) when compared to the native protein (C121). The phases were solved using the SAD phasing options in CRANK2^15^ without using structural information from AlphaFold2. The model obtained from CRANK2 had an *R*_work_ and *R*_free_ of 0.35 and 0.42, respectively. The output from CRANK2 was used as a search model for native OmamC_129-261_ data in PHASER which provided a solution with a TFZ and LLG scores of 21.5 and 742, significantly higher than the values obtained from the AlphaFold2 predicted structure. Further modelbuilding by COOT^16^ and refinement in REFMAC5^17^ and PHENIX^18^ improved the model to final *R*_work_ and *R*_free_ of 0.228 and 0.253. Although native and SeMet-labelled OmamC_129-261_ crystallized in different space groups, both have three molecules in the asymmetric unit and are in the same arrangement forming a funnel shaped trimer with the N-terminal end of the helix close to each other (**Fig. 3A and B**).

**Fig 3.**
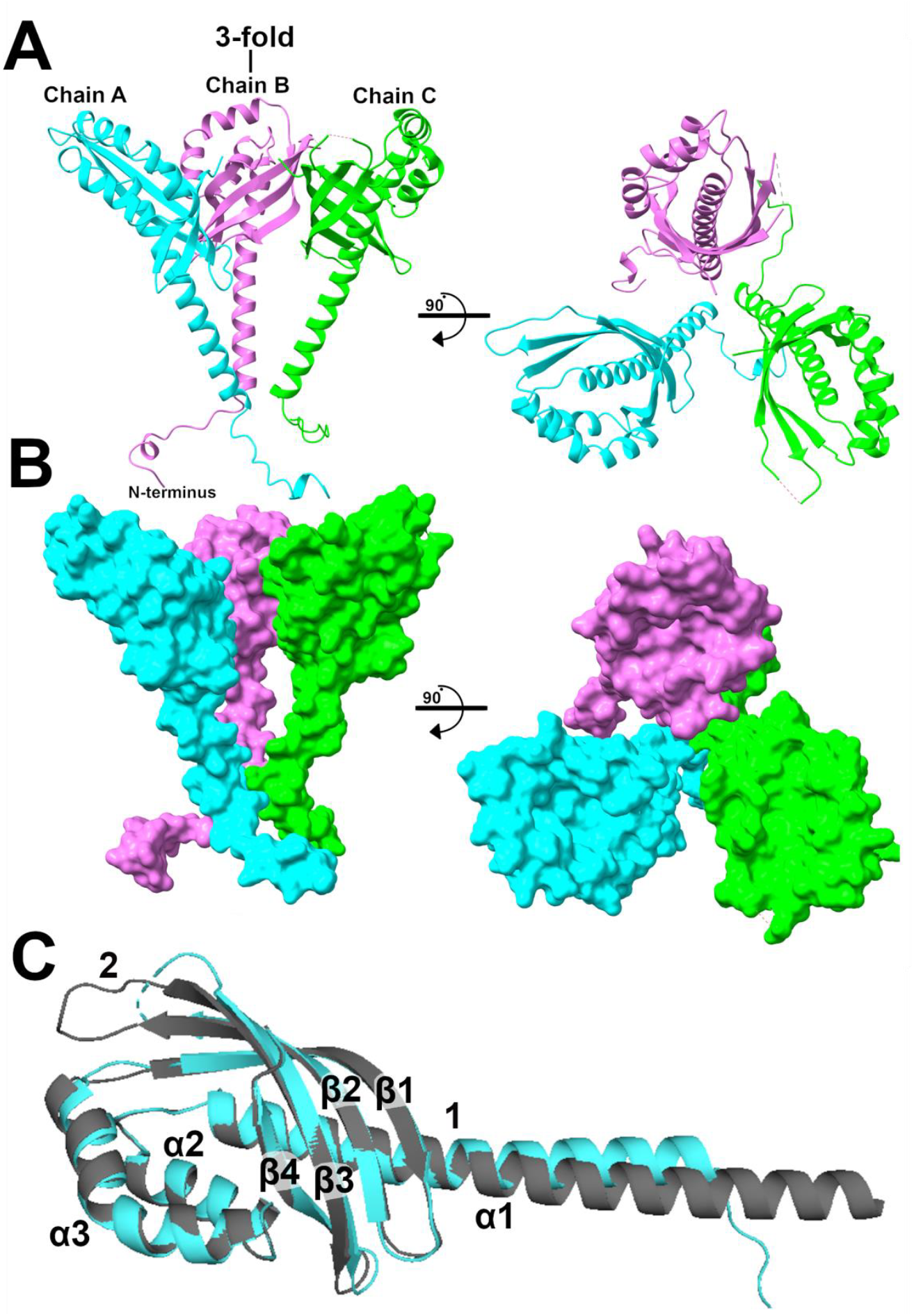
Trimer of OmamC_129-261_ as observed in the asymmetric unit of the crystal structure with a funnel like arrangement shown in side and top view in **A)** cartoon representation and **B)** surface representation. **C)** Superposition of chain A of OmamC_129-261_ crystal structure (cyan) to the AlphaFold2 predicted structure (black). The key differences between the predicted and experimentally determined structure are highlighted as 1 and 2.

When comparing the monomeric structure of OmamC_129-261_ with that of the AlphaFold2 predicted structure, the overall topology is similar. On a closer look, in the crystal structure of OmamC_129-261_, the α1-helix has a bent of about 20° at A141 whereas such a bend was not present in the AlphaFold2 predicted structure **(Fig. 3C, label 1)**. The α3-helix has also bent by about 60° at A186, which has been correctly predicted in the AlphaFold2 predicted structure (**Fig. 3C**). Noteworthy, both the α helical bends have a sequence motif of QQRAA and the bend occurs after the last Ala. Other differences between the predicted and crystal structure were noted in the loop region, where some of the loop region connecting the β-strands had differences. The loop region between β2 and β3 helices had very poor electron density in the crystal packing, which could be caused by a flexibility in this region (**Fig. 3C, label 2**). Missing residues in shown as dotted line. These small but critical differences seem to have a major impact on the structure determination using the AlphaFold2 predicted model, emphasizing the importance of experimental structure determination.

## Concluding remarks

During the crystallization and structure determination process of OmamC_129-261_, several challenges were encountered, and interesting observations were made. In the beginning, the use of a specific imine solution was critical. Alternative reservoir method also produced reproducible and big crystals, although their diffraction quality was very poor. However, better quality in diffraction and reproducibility were obtained by removing the partially folded proteins by a short heat treatment. Another interesting observation was that despite the overall similarity, the predicted structure was not sufficiently accurate to determine the structure by molecular replacement alone. The heat treatment to temperatures close to onset of unfolding temperature of the protein has played a critical role in obtaining good quality crystals OmamC_129-261SeMet_ crystals, which ultimately led to the structure determination. Noteworthy, this is the first experimentally determined structure of any Mam/Omam proteins. Although both the heat-treatment and alternative reservoir methods have been reported earlier, they are apparently not as widely used methods. It is worthy to try these methods, especially when there are challenges with the reproducibility and obtaining diffraction quality crystals.

## Methods

### Cloning, expression and purification

*omamC* (*rv1363c*) was amplified from Mtb genomic DNA and inserted into to the multiple cloning site (MCS) of the petM11 vector in a restriction-free cloning approach. Similarly, a construct for expressing truncated variant of OmamC (OmamC_129-261_) was generated in which the variable N-terminal region and membrane spanning helix are lacking. The correctness of the inserted sequences was confirmed by DNA sequencing at the Biocenter Oulu sequencing core facility. The OmamC and OmamC_129-261_ constructs were separately transformed into *E. coli* Bl21DE3 strains and cultivated at 37 °C in LB media containing appropriate antibiotic. The overexpression of protein was induced with 0.4 mM β-D-1-thiogalactopyranoside (IPTG), once OD_600_ of 0.6-0.9 was reached and incubated overnight for expression at 20 °C. Harvested cells were resuspended in a lysis buffer (50 mM Tris, 500 mM NaCl, 2.5 mM DTT at pH 8.5) with added 20 µg ml^-1^ of DNase and RNase, 0.1 mg ml^-1^ lyzozyme, and 5 mM MgCl_2_and lysed by passing the suspension through cell disruptor at 34.8 kPSi twice. The lysed solution was then left rotating at 6 °C for an hour. Cell debris was removed by centrifugation at 30 000 rcf for 45 min at 6 °C. OmamC_129-261_ was further purified with IMAC using Ni-NTA beads, supplementing the lysis buffer with 50 mM of imidazole for washing and 500 mM for eluting. The eluted sample was directly injected into HiLoad 16/60 Superdex 200 prep grade 120 ml SEC column, which was pre-equilibrated with a 50 mM MOPS buffer containing 300 mM KCl and 2.5 mM DTT at pH 7.0. Pure fractions were analysed by SDS-page, concentrated, flash frozen in liquid nitrogen and stored at -70 °C. SeMet-labelled variant (OmamC_129-261SeMet_) expression was done using minimal media with supplemented SeMet to replace native methionine in the protein complex. Expression and purification were carried out the same way as unlabelled OmamC_129-261_ variant.

### Thermal stability and folding

For secondary structure analysis, protein was diluted to 0.1 mg ml^-1^ in 10 mM potassium phosphate buffer at pH 7.6. Instance of aggregation as well as *T*_m_ was measured by nanoDSF at concentration of 0.1 mg ml^-1^, 0.3 mg ml^-1^, and 0.6 mg ml^-1^ from 20 °C to 90 °C with 1 °C per minute temperature increase. The nanoDSF results were analysed using MoltenProt^19^ webservice, in which data quality and *T*_m_ was calculated.

### Crystallization, data collection and structure determination

Diffraction quality crystals for both native and SeMet-labelled OmamC_129-261_ were obtained (3-6 mg ml^-1^ in 50mM MOPS, 300mM KCl, 2.5mM DTT at pH 7.0) in sitting drop vapor diffusion method at 6 °C in 2 to 1 ratio of protein to reservoir solution of the MIDASplus screen H03 sub-well condition containing 100 mM Tris with 20% glycerol ethoxylate and 3% imine polymer at pH 8.5. Progress of crystallization was monitored online with Icebear software^20^. Nucleation of crystals was observed within 24 hours. OmamC_129-261_ was heat treated at 35 °C for 15 minutes followed by spinning the sample at 15 000 rcf for 10 minutes at room temperature (RT). Supernatant was carefully removed to a precooled Eppendorf tube and kept on ice for 30 minutes. If appearance of cloudiness or visible formation of aggregation was observed, the sample was centrifuged again as above or filtered through 0.2 µm filter. After the heat treatment, the final concentration of the protein was reduced by 30 to 50 % of the starting protein concentration and crystallization performed in as described above. In addition, crystallization trials were also conducted in alternative reservoir method^14^, to discover new nucleation conditions. The desired precipitant solution was mixed with protein solution as with normal crystallization plate, but reservoir solution was replaced with 1.5 M NaCl.

X-ray diffraction data was collected for OmamC_129-261_ and OmamC_129-261SeMet_ at the I04 beamline of Diamond Light Source, Didcot, UK to a resolution of 2.3 and 3.14 Å, respectively. Data was processed with xia2.dials^21,22^ and analyzed with AIMLESS in CCP4i2 cloud service^23^. Initial phases were calculated by SeMet-SAD phasing from the data collected from OmamC_129-261SeMet_ crystals with CRANK2 pipeline^15,24^. Partially build model obtained from this step was used for molecular replacement in PHASER^25^. The resulting model was improved further with iterative cycles of model building in COOT^16^ and refinement in REFMAC5^17^ from CCP4i2 cloud services and phenix.refine from the Phenix software package^16–18,25,26^.

## Acknowledgement

We acknowledge Dr. Sven Sowa for suggesting heat treatment to improve crystallization. We acknowledge Biocenter Oulu structural biology, proteomics and protein analysis, and DNA sequencing core facilities for their support. We acknowledge the funding support from the Tampere Tuberculosis Foundation, Research Council of Finland (332 967 & 371 075) and the University of Oulu Graduate School.

## Data availability

The relevant data associated with the published study are present in the paper or attached as Supplementary data. Structural data is submitted to the PDB with the accession code pdb_00009R33.

## Author contributions

R.V. with M.J.H. planned the study, analyzed the results and wrote the manuscript. M.J.H. performed all the experiments. R.V. acquired the funding for the work.

